# Testing the ecological and behavioural responses of mammals to the Landscape of Fear

**DOI:** 10.1101/2025.06.05.657929

**Authors:** Stefano Tolusso, Giovanni Zanfei, Alessio Mortelliti

## Abstract

The fear of being predated can influence many decisions taken by preys and this may have important consequences for the ecosystems. Preys can respond to the risk of being predated in three ways: habitat selection, diel activity modifications and adoption of behaviors to detect predators before they constitute a danger. However, rarely these strategies have been analyzed simultaneously within an entire community. Through a camera trapping study conducted in the Julian Prealps Natural Park and surrounding territories (North-East of Italy), we analyze these three antipredator responses to understand which strategy is adopted by target species. Results show how, for the majority of species, the main antipredator response is the selection of time intervals when corresponding predators are inactive. On the opposite, habitat selection seems to be more influenced by environmental characteristics of the area. Similar, behavioral traits do not result related to presence probability of predators. Therefore, in the area analyzed, we demonstrated the existence of a landscape of fear that influences species choices regarding mainly diel activities. The study underlines also the importance of simultaneous consideration of antipredator responses for a more accurate understanding of how fear structures communities and ecosystems.

## INTRODUCTION

By experiencing the emotion of fear, animals are able to understand when their life is at risk, which is particularly important for preys in the presence of predators. Fear is associated with a perception of risk, which varies in space as some areas are perceived being more risky than others, and this spatial variation is known as the Landscape of Fear, which affects multiple aspects of the ecology of a species (Gaynor et al., 2019). Risk perception may depend upon different factors: such as body condition as starving individuals may decide to take higher risks compared to individuals with better body condition (Bleicher, 2017). Other factors affecting risk perception are age and sex (Burton et al., 2022; Hunter and Skinner, 1998), personality (Boone et al., 2022; Heithaus et al., 2007; Merz et al., 2023) and sociality (Bleicher, 2017). Risk perception and thus the landscape of fear, may also vary depending on the type of predators present, including their hunting strategies (i.e. cursorial vs stalker) (Chitwood et al., 2022; Epperly et al., 2021; Gallagher et al., 2017; Thaker et al., 2011). Based on this spatial distribution of risk, animals take decisions, facing a tradeoff between food acquisition and safety (Gaynor et al., 2019). However these decisions may vary with time, mainly influenced by resource abundance (Riginos, 2015)

Preys respond to risk through three non-mutually exclusive strategies: a) *habitat selection*: they attempt to avoid areas where the predator is present, b) *activity patterns*: they concentrate activities in times of the day when predators are less active and c) *antipredatory behaviors*: they display behaviors such as vigilance, to maximize the detection of danger (Brown, 1999; Creel and Christianson, 2008; Dröge et al., 2017). The strategy adopted will affect the fitness of an individual (Mitchell and Harborne, 2020; Zanette and Clinchy, 2020), but also the whole community and ecosystems, because avoidance of certain areas by preys can determine resource release for other species, and thus have cascade effects on the trophic chain (Chitwood et al., 2022; Laundré et al., 2001; Ripple and Beschta, 2004). Fear in mesopredators can have positive consequences for their preys that usually respond by increasing in number (Newsome et al., 2017).

The way preys cope with the perception of risk may have important consequences for conservation and management. Changes in habitat structure may contribute to strengthen antipredator responses or alter predators hunting efficiency (Gaynor et al., 2021; Sandford et al., 2017). Similarly, introduction of risk cues (i.e. cues that may indicate the presence of a predator in a certain area) may push species away from certain areas preserving endangered plant species or reducing human-wildlife conflicts (Chitwood et al., 2022; Gaynor et al., 2021; Ramirez et al., 2024).

Despite the growing interest on this topic, most studies focus only on one of the possible antipredator responses, usually with contrasting results. Indeed, it is often difficult to demonstrate the adoption of a particular antipredator strategy (Kauffman et al., 2010; Kohl et al., 2019; Schmidt and Kuijper, 2015). We are aware of only one study considering all three strategies in mammalian species (Muthersbaugh et al., 2025). Identifying the most commonly adopted approach, which is only possible if all three strategies are considered together, will help shed the light on mechanisms facilitating predator-prey coexistence, which is particularly important in modified landscapes where shifts in habitat selection or activity patterns may not be feasible, or where vigilance behavior may not be adapted to the arrival of new predator species (Kohl et al., 2018; Lone et al., 2014). When all responses are considered it is possible to make more accurate inferences regarding which type of cues are related to preys risk perception, or at which scale different choices of animal took place. Our goal here is to contribute in filling this knowledge gap on the relative importance of the three strategies to cope with the Landscape of fear. Thus, our objectives are to test the relative importance of the three strategies adopted by preys, specifically a) if and to what extent predators’ presence influences preys habitat selection b) if preys concentrate activity during time intervals in which predators are less active and c) if the presence of predators determines the adoption of behavioral traits useful to detect the presence of a danger. To achieve our objective we conducted a field study in the Italian Alps, in an area encompassing multiple habitat types including beech forest, conifer forest, prairies and riparian habitats. We considered a large part of the mammalian community, including carnivores such

as the golden jackal (*Canis aureus*), fox (*Vulpes vulpes*), badger (*Meles meles*) and stone marten (*Martes foina*), artyodactyla: red deer (*Cervus elaphus*), roe deer (*Capreolus capreolus*), chamois (*Rupicapra rupicapra*) and wild boar (*Sus scrofa*) and small mammals (*Apodemus flavicollis, A. sylvaticus, A. agrarius*), dormouse (*Glis glis*) and squirrel (*Sciurus vulgaris*). The wide taxonomic scope of our study thus allowed us to consider multiple predator–prey interactions such as between ungulates and golden jackal, or small mammals and canids, or between carnivores of different sizes.

## MATERIALS AND METHODS

### Study area

The study was conducted in the Julian Prealps Natural Park and surrounding areas in the Friuli Venezia Giulia region (Italy). The study area included the pre-Alpine mountain belt with the highest mountains reaching 1900 m of altitude, except for “Monte Canin” (2587 m). The area is characterized by relatively high rainfall, with more than 3300 mm of rain each year (ARPA FVG 2023). The mean annual temperature is approximately 12° in the valley floor. The area is dominated by beech forests (*Fagus sylvatica*), coniferous forests dominated by pines (*Pinus nigra* and *Pinu sylvatica*) typically growing in areas following landslides or fires. The upper part of the mountains are dominated by prairies *Sesleria cerulea* and *Carex sempervirens* more or less invaded by different species of shrubs and bushes.

#### Camera trapping

We conducted our camera trapping survey in 2024 from 14 June to 3 September. We surveyed a total of 64 sites (*sensu* Mackenzie et al., 2002) stratified by the four habitat types: beech forests, coniferous forests, alpine prairies and riparian habitats, with 16 sites in each habitat. During each sampling session we simultaneously sampled 4 sites in each habitat type. Sites were spaced >1km apart to maximize independence (i.e. lower the chances that the same individual would visit >1 site). Each site was composed by two camera stations distanced 100 m (Evans et al., 2019). In one of the stations we used a can of sardines as attractant; we elected to use sardines as they are the most cost-effective attractant for the target mammalian community (Mortelliti et al., 2024). The can was perforated in the upper part to spread scent and the content was thus not accessible to animals (hence it worked as a lure, not as bait (Buyaskas et al., 2020; Mortelliti et al., 2024)). Camera traps (*Browning Recon Force Elite* model *BTC-7E-HP5*) were positioned 30/40 cm above ground and set to take 20 s videos when an animal cross their field of view (trigger speed 0.1 s), with a delay between each video of 1 minute. Each site was active from a minimum of 14 to a maximum of 20 days.

#### Small mammal trapping

Small mammals’ presence and abundance data was collected in September and October, following the same camera trapping design. We sampled each site with a linear transect of 10 “Heslinga” traps spaced 10 m apart activated for 3 consecutive nights. Traps were baited with a mixture of flour, seeds, peanut butter and water and cotton was provided for bedding. The small mammals captured were marked trough haircut and we took a sample of the ear for genetic confirmation of species. In addition, we collected the following environmental variables for each transect: shrub cover at height of 2 and 6 m above ground, forest cover (in a scale from 1 to 4) and presence of rocks (1 to 3), all variables we thought may potentially influence the presence of small mammals in the area. We than calculated the Minimum Number Known Alive (MNKA), which is the minimum number of individuals captured along the transect. Hazel dormice presence/absence data were collected using nest-tube transects with 5 nest tubes (67 x 67 x 297 mm) spaced 20 m apart. We positioned 30 tubes per habitat type in beech forests, pine forests and shrubby areas for a total of 60 transects (30 of which were positioned in the same sites as the camera traps). Environmental data was collected through a GIS, using Corine biotopes classification obtained in a 500 m buffer around the center of each camera trap site created in QGIS 3.16.11. The areas were grouped into the following macro-categories: beech and coniferous forests, open, urban and riparian areas. The surfaces were then related to the total buffer area. We also calculated the distance of each site from the nearest urban area (both a small village and industrial areas). Additional data derived from Digital Terrain Model (DTM) of the area from which we calculate altitude and slope of each site (for the slope we retain the median value in each buffer). When considered meaningful for the species under analysis (i.e. dormouse and squirrel) we calculated 100 and 200 buffers around the center of each camera trap site.

#### Behavioral traits measurement

To quantify risk perception we measured a set of behavioral traits, depending on taxa. For ungulates, we measured 1) *vigilance levels*: measured as the proportion of time an animal spent with its head above the shoulders line, with the ears raised and not chewing (Hunter and Skinner, 1998; Le Saout et al., 2015); 2) *number of flights:* each time an animal crossed the camera field of view at high speed, when it escaped during the video, such as at the activation of the camera trap. Both traits were related to the time individuals of each video remains in the camera field of view in each site. If the head of an animal was not visible, this amount of time was not considered. For the other species we considered the interaction with the attractant; the behaviors considered were: 1) *latency at the contact* 2) *indifference* 3) *duration of interaction with the attractants* (Del Fabbro et al., 2023, unpublished data). These behaviors were considered as proportions of the time each individual spent in in the camera field of view. The first two traits were considered as indicators of an animal’s perception of high risk, whereas the conversely for the interaction.

### Statistical analysis

We performed three main statistical analyses in order to understand how fear influenced species decisions and behaviors. We restricted our analyses to species present in more than 10% of sampled sites (Mortelliti et al., 2022).

#### Occupancy modelling

First, we fitted single season *occupancy* models to detection history data to identify the main predictors of species’ distribution and detection in the area sampled. We considered a “visit” to each site as a 24 hours period starting noon of the first day camera displacement (Evans and Mortelliti, 2022). We used the following covariates as predictors for Ψ (presence probability): presence probability of prey/predator, percentage of habitat, slope, elevation, MNKA, habitat typology (categorical) and shrub and tree cover. For p (detection probability): percentage of habitat, time since camera deployment and precipitations. We followed a forward stepwise approach to model selection, therefore we started by modelling detection probability, with presence held constant; the top ranking model was then retained and we repeated the process for presence probability. We calculated the overdispersion parameter (*c-hat*) and we ranked models by AICc (i.e. corrected for small sample size) using all models within 2 ΔAICc for inference (Anderson and Burnham, 2002). Since the red squirrel and the stone marten were never detected in prairies we exclude these sites from the analyses.

#### Analysis of activity patterns

We conducted an overlap analysis (Meredith and Ridout, 2014) to test the hypothesis that preys were active during intervals of time in which their predators are less active. We first calculated daily activity patterns, obtained by kernel density estimations, of all single species and we then obtained overlap coefficients (Δ_1_) (Meredith and Ridout, 2014). The predators considered for comparison were golden jackal, fox and stone marten. For the coefficient calculation we exclude prairies sites due to lower predators presence in these areas and we also split red deer dataset into adults and fawns because of the vulnerability of the latter to golden jackal predation. We used the Watson two sample test for circularity to quantify differences between activity patterns of species (Pewsey et al., 2013). We then calculated confidence intervals for each overlap coefficient between preys and predators with 10000 bootstraps (Meredith and Ridout, 2014). The intervals were then compared among each other.

#### Analysis of behavioral traits

We use Generalized Linear Mixed Models (GLMM) to understand if the presence probability of predators may affect time spent displaying vigilance behaviors. The site ID was used as random factor to account for the lack of independence of repeated observations. In the case of roe deer we added an additional categorical variable to take into account the mating period (which would affect certain behaviours such as the proportion of time spent in marking territory and other mating behaviors), for red deer we included a binary variable to account for the presence of fawns (due to their vulnerability to predation). We selected the binomial distribution, with logit link function; models were ranked based on AICc. Before the analysis all variables were checked for collinearity and standardized. We used R 4.3.1 statistical software and *unmarked*, *overlap* and *lme4* packages (Bates et al., 2003; Fiske and Chandler, 2011; Meredith and Ridout, 2014).

## RESULTS

Our camera-trapping survey encompassed a total of 1790 trap nights; the most widespread species was red deer, found in 49 of 64 sites (76.6%), followed by the roe deer which was found in 42 (65.6%) sites. We then captured the red fox in 32 sites (50.0%), the stone marten in 24 (37.5%), the golden jackal in 18 (28.1%), and the chamois in 13 (20.3%). The European squirrel and the fat dormouse were detected 11 sites (17.2%), whereas the wild boar and the badger were found in 10 sites (15.6%). We also found in a limited number of sites the brown hare (*Lepus europaeus*) (10.9%) and the European polecat (*Mustela putorius*)(3.1%). The pine marten (*Martes martes*), the wild cat (*Felis sylvestris*), the otter (*Lutra lutra*) and the black rat (*Rattus rattus*) were found in only one site. We captured a total of 124 small mammals (out of 41 transects,) including 102 *Apodemus flavicollis,* 6 *Glis glis,* 4 *Sorex araneus,* 3 *Apodemus agrarius,* 3 *Myodes glareolus,* 2 *Sorex alpinus,* 2 *Crocidura leucodon,* 1 *Crocidura suaveolens* and 1 *Suncus etruscus.* We found the hazel dormouse in 5 sites.

### Occupancy analysis

The probability of presence of predators was not included in the top ranking models for any of the analyzed prey species. Only the second-ranked model for the red fox included a negative relationship with the probability of presence of the golden jackal. On the opposite, we found a positive relationship between the probability of presence of the golden jackal and the roe deer (Fig. 3b).

Among ungulates detection probability, the red deer shows a decreasing relation with an increasing proportion of riparian habitat (Fig. 2a) and the chamois has the same relation with coniferous forest abundance (Fig. 2c). On the opposite, wild boar detection probability increases with beech forest proportion (Fig. 2d). Only for roe deer, the coefficient estimation result is constant: p _ROE DEER_ = 0.23 (+/- 0.03). On the other side, presence probability of roe deer decreases as slope increases (Fig. 2b), of wild boar, increases with the abundance of urban areas (Fig. 2e) and decreases with increasing proportion of beech forests (Fig. 2f). The presence probability of red deer and chamois result constant among sites in the studied area: Ψ_RED DEER_ = 0.82 (+/- 0.08) and Ψ_CHAMOIS_ = 0.30 (+/- 0.11). Golden jackal presence probability does not result in affecting prey ones. In table 1 models within 2 ΔAICc for ungulate species of the studied area.

**Figure 1:**
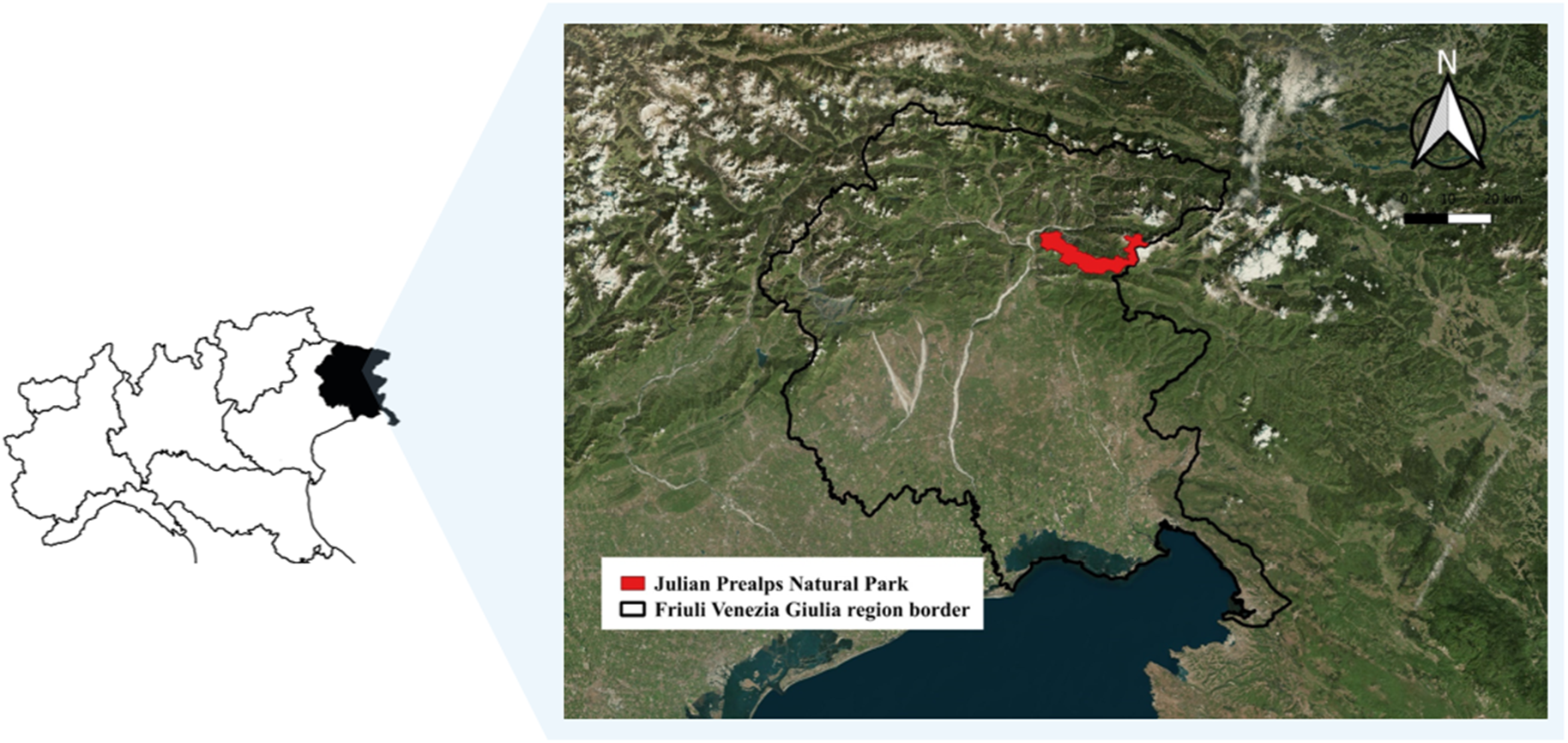
Location of the study area in the Julian Prealps Natural Park, Friuli Venezia Giulia region, Italy.

**Figure 2:**
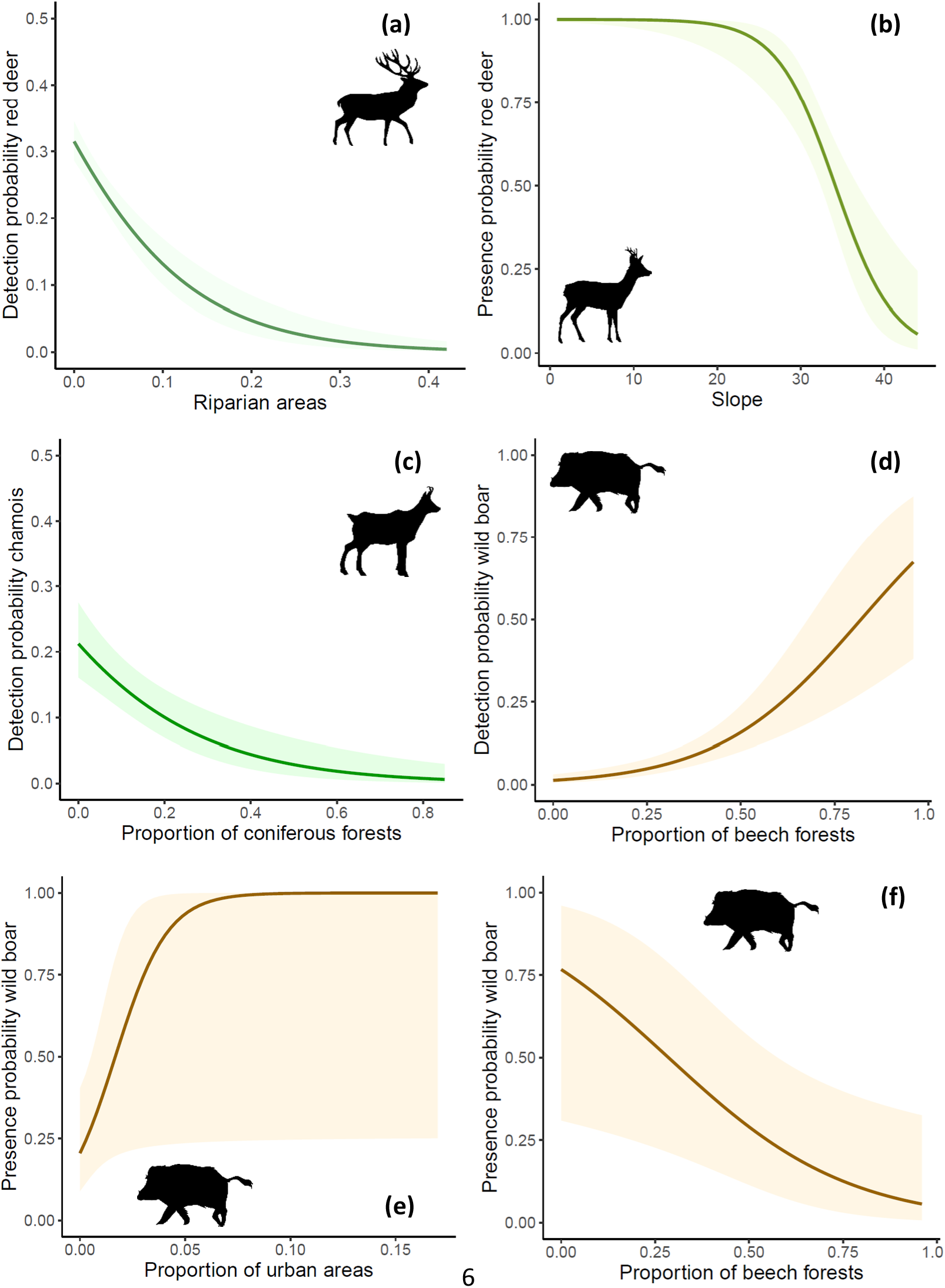
predictions of the top ranked single season models for ungulates. Site is intended as 500 m buffer around transect central point. The 85% confidence interval is shown with lighter colors. a) red deer detection probability decrease with the proportion of riparian areas b) roe deer presence probability decreases with increasing slope of the terrain; c) chamois detection probability decreases with coniferous forest abundance; d) wild boar detection probability increases with the proportion of beech forests in the landscape; e) and f) wild boar presence probability increases with the percent cover of urban areas and decreases with the proportion of beech forest in the landscape.

**Table 1:** Top ranked (within 2 ΔAICc) single season occupancy models for ungulate species; p = detection probability, Ψ = presence probability of considered species. Selected variables are in brackets, habitat types are intended as proportion in each site. Point in brackets is intended absence of predictors. For each model we report: distance from top ranking model (ΔAICc), AICc value, number of parameters (Par) and R^2^ (Rsq).

| Occupancy models ungulates |  |  |  |  |  |
| --- | --- | --- | --- | --- | --- |
| Species | Model | $\Delta AICc$ | AICc | Par | Rsqu |
| Red deer | $p(\text{riparian areas})\Psi(\text{slope})$ | 0 | 887.52 | 4 | 0.41 |
| | $p(\text{riparian areas})\Psi(.)$ | 0.16 | 887.68 | 3 | 0.38 |
| | $p(\text{riparian areas})\Psi(\text{riparian areas})$ | 1.07 | 888.59 | 4 | 0.42 |
| | $p(\text{riparian areas})\Psi(\text{slope} + \text{riparian areas})$ | 1.11 | 888.63 | 5 | 0.40 |
| | $p(\text{riparian areas})\Psi(\text{beech forests})$ | 1.81 | 889.33 | 4 | 0.39 |
| Roe deer | $p(.)\Psi(\text{slope})$ | 0 | 753.44 | 3 | 0.22 |
| Chamois | $p(\text{coniferous forests})\Psi(\text{wood})$ | 0 | 233.84 | 4 | 0.23 |
| | $p(\text{coniferous forests})\Psi(\text{slope})$ | 0.20 | 234.04 | 4 | 0.21 |
| | $p(\text{coniferous forests})\Psi(\text{cliffs})$ | 0.22 | 234.05 | 4 | 0.20 |
| | $p(\text{coniferous forests})\Psi(\text{wood} + \text{slope})$ | 0.29 | 234.13 | 5 | 0.20 |
| | $p(\text{coniferous forests})\Psi(\text{wood} + \text{cliffs})$ | 0.94 | 234.78 | 5 | 0.22 |
| | $p(\text{coniferous forests})\Psi(\text{beech forests})$ | 0.97 | 234.81 | 4 | 0.19 |
| | $p(\text{coniferous forests})\Psi(.)$ | 1.02 | 234.86 | 3 | 0.19 |
| | $p(\text{coniferous forests})\Psi(\text{open areas})$ | 1.13 | 234.97 | 4 | 0.14 |
| | $p(\text{coniferous forests})\Psi(\text{altitude})$ | 1.75 | 235.59 | 4 | 0.24 |
| Wild boar | $p(\text{beech forests})\Psi(\text{urban areas} + \text{beech forests})$ | 0 | 143.74 | 5 | 0.28 |
| | $p(\text{beech forests})\Psi(\text{urban areas})$ | 1.88 | 145.62 | 4 | 0.22 |

Among carnivore detection probabilities, the golden jackal shows a decreasing relation with the increase of the distance from urban areas (Fig. 3a); the fox shows an increasing relation with the abundance of human-inhabited areas (Fig. 3d) and a parabolic negative relation with time since camera activation, with the highest detection probability around the 7^th^ day (Fig. 3e). The detection probability of stone marten increases with open areas abundance (Fig. 3g) and decreases with riparian habitats percentage (Fig. 3h), for badger, the value is constant across the studied area: p_BADGER_ = 0.11 (+/- 0.04). Jackal presence probability decreases with altitude (Fig. 3c) and increases with roe deer presence probability (Fig. 3b). Fox presence probability is higher in sites containing urban areas (Fig. 3f); however, in the second-ranked model, the presence probability of jackal negatively affects fox one. Stone marten presence probability increases with abundance of urban areas (Fig. 3j) and is higher in riparian habitats while lowest values are recorded in beech forests (Fig. 3i). For badger, also presence probability results constant: Ψ_BADGER_ = 0.19 (+/- 0.09). In table 2 models within 2 ΔAICc for carnivores species of the studied area.

**Figure 3:**
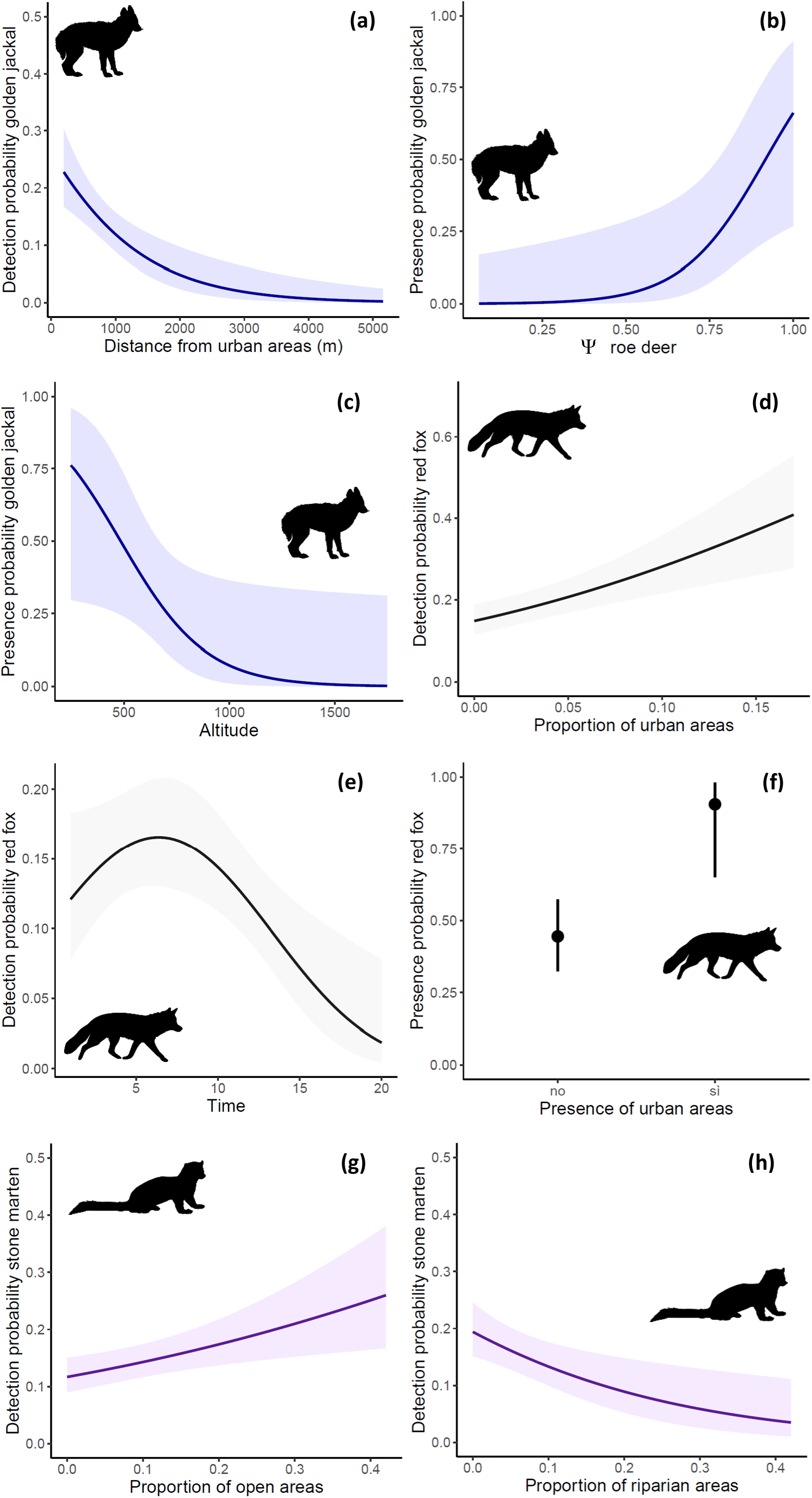
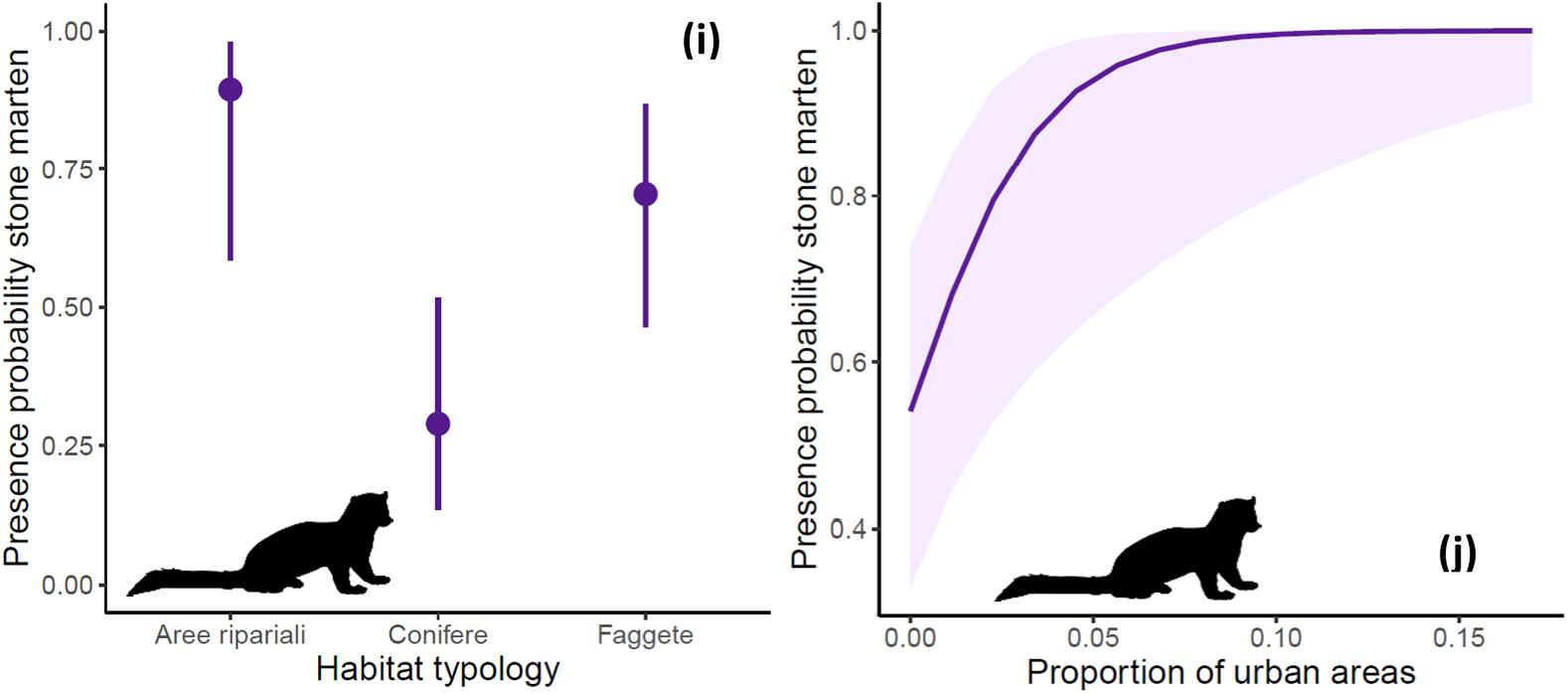
single season models previsions for carnivores. Site is intended as 500 m buffer around transect central point. The 85% confidence interval is shown with lighter colors. In a) golden jackal detection probability decrease with distance from urban areas; b) and c) golden jackal presence probability increase with roe deer presence probability and decrease with altitude; d) and e) fox detection probability increase with urban areas abundance and follow a parabolic trajectory over time; f) fox presence probability is higher when urban areas are present inside sites; g) and h) stone marten detection probability increase with open areas percentage and decrease with riparian areas abundance; i) and j) stone marten presence probability is higher in riparian areas and lower in beech forests and increases with urban areas percentage.

**Table 2:** Top ranked (within 2 ΔAICc) single season occupancy models for carnivore species; p = detection probability, Ψ = presence probability of considered species. Selected variables are in brackets, habitat types are intended as proportion in each site. Point in brackets is intended absence of predictors. For each model we report: distance from top ranking model (ΔAICc), AICc value, number of parameters (Par) and R^2^ (Rsq).

| Occupancy models carnivores |  |  |  |  |  |
| --- | --- | --- | --- | --- | --- |
| Species | Model | $\Delta AICc$ | $AICc$ | Par | Rsqr |
| Golden jackal | $p(\text{cities distance})\Psi(\text{altitude} + \Psi \text{ roe deer})$ | 0 | 292.21 | 5 | 0.50 |
| | $p(\text{cities distance})\Psi(\text{altitude} + \Psi \text{ roe deer} + \Psi \text{ Apodemus})$ | 1.11 | 293.32 | 6 | 0.51 |
| | $p(\text{cities distance})\Psi(\Psi \text{ Apodemus} + \Psi \text{ roe deer})$ | 1.16 | 293.37 | 5 | 0.49 |
| | $p(\text{cities distance})\Psi(\text{altitude} + \text{slope})$ | 1.22 | 293.43 | 5 | 0.49 |
| | $p(\text{cities distance})\Psi(\Psi \text{ roe deer})$ | 1.46 | 293.67 | 4 | 0.47 |
| Fox | $p(\text{urban areas} + \text{time}^2)\Psi(\text{urban p/a})$ | 0 | 505.59 | 6 | 0.31 |
| | $p(\text{urban areas} + \text{time}^2)\Psi(\text{urban p/a} + \Psi \text{ golden jackal})$ | 1.73 | 507.32 | 7 | 0.32 |
| Stone marten | $p(\text{open areas} + \text{riparian areas})\Psi(\text{urban areas} + \text{habitat typology})$ | 0 | 385.51 | 7 | 0.35 |
| Badger | $p(\text{open areas})\Psi(\text{urban areas})$ | 0 | 167.12 | 4 | 0.10 |
| | $p(.)\Psi(\text{urban areas})$ | 0.51 | 167.64 | 3 | 0.06 |
| | $p(\text{open areas})\Psi(\text{beech forests})$ | 0.58 | 167.70 | 4 | 0.09 |
| | $p(.)\Psi(\text{altitude})$ | 0.71 | 167.83 | 3 | 0.05 |
| | $p(\text{open areas})\Psi(\text{altitude})$ | 1.05 | 168.17 | 4 | 0.08 |
| | $p(\text{open areas})\Psi(\text{wood})$ | 1.16 | 168.29 | 4 | 0.08 |
| | $p(\text{open areas})\Psi(\text{coniferous forests})$ | 1.36 | 168.48 | 4 | 0.08 |
| | $p(.)\Psi(\text{coniferous forests})$ | 1.41 | 168.53 | 3 | 0.04 |
| | $p(\text{open areas})\Psi(\Psi \text{ sciacallo})$ | 1.68 | 168.80 | 4 | 0.07 |
| | $p(.)\Psi(.)$ | 1.77 | 168.90 | 2 | 0 |

The last species analyzed are two arboreal rodents: the fat dormouse and the squirrel. Dormouse detection probability increases with the percentage of woods (conifer, beech and mixed forests) within a buffer of 100 m (Fig. 4a), while that of the squirrel increases with the proportion of coniferous forests of the site (Fig. 4b). In addition, the latter detection probability varies with time since transect activation with higher detectability values at the beginning and at the end of sampling period (Fig. 4c). The presence probability results constant for both species: Ψ_DORMOUSE_ = 0.39 (+/- 0.17), Ψ_SQUIRREL_ = 0.38 (+/- 0.16); however in either case, in models within 2 ΔAICc, is included stone marten presence probability with positive relation with dormouse and negative with squirrel presence probability. In table 3 models within 2 ΔAICc for arboreal rodent species of the studied area.

**Figure 4:**
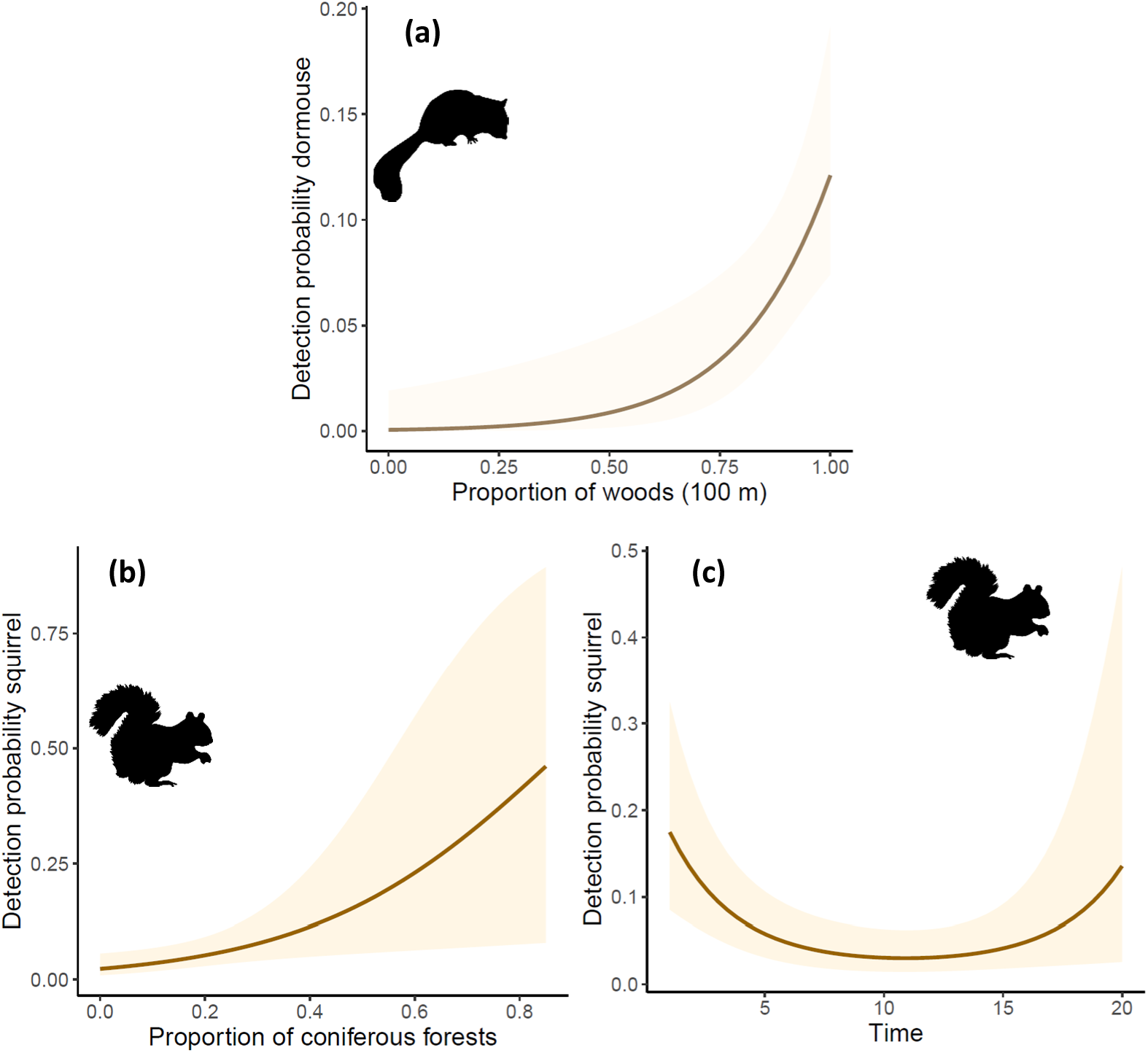
single season models previsions for rodents. Site is intended as 500 m buffer around transect central point or when specified 100 m. The 85% confidence interval is shown with lighter colors. In a) dormouse detection probability increases with wood percentage in 100m radius; b) and c) squirrel detection probability increases with coniferous forests abundance and is higher at the beginning and at the ending of sampling period

**Table 3:** Top ranked (within 2 ΔAICc) single season occupancy models for rodent species; p = detection probability, Ψ = presence probability of considered species. Selected variables are in brackets, habitat types are intended as proportion in each site. Point in brackets is intended absence of predictors. For each model we report: distance from top ranking model (ΔAICc), AICc value, number of parameters (Par) and R^2^ (Rsq).

| Occupancy models rodents |  |  |  |  |  |
| --- | --- | --- | --- | --- | --- |
| Specie | Modello | $\Delta AICc$ | AICc | Par | Rsqr |
| *Fat dormouse* | $p(\text{wood } 100)\Psi(\Psi \text{ stone marten})$ | 0 | 165.30 | 4 | 0.18 |
| | $p(\text{wood } 100)\Psi(.)$ | 0.60 | 165.90 | 3 | 0.12 |
| | $p(\text{wood } 100)\Psi(\text{wood } 200)$ | 0.71 | 166.01 | 4 | 0.16 |
| | $p(\text{wood } 100)\Psi(\text{open areas } 200)$ | 1.06 | 166.35 | 4 | 0.16 |
| | $p(\text{wood } 100)\Psi(\Psi \text{ fox})$ | 1.95 | 167.24 | 4 | 0.14 |
| | $p(\text{wood } 100)\Psi(\text{bushes cover at } 6 \text{ m})$ | 1.95 | 167.25 | 4 | 0.14 |
| *Squirrel* | $p(\text{coniferous forests} + \text{time}^2)\Psi(\text{habitat typology})$ | 0 | 166.44 | 7 | 0.37 |
| | $p(\text{coniferous forests} + \text{time}^2)\Psi(\Psi \text{ stone marten})$ | 1.03 | 167.48 | 6 | 0.32 |
| | $p(\text{coniferous forests} + \text{time}^2)\Psi(.)$ | 1.73 | 168.18 | 5 | 0.26 |

### Overlap analysis

The majority of species shows crepuscular or nocturnal activity patterns with the exception of the squirrel, which results being diurnal, and the chamois that shows, in addition to crepuscular activity, high activity during the day (Fig. 5).

**Figure 5:**
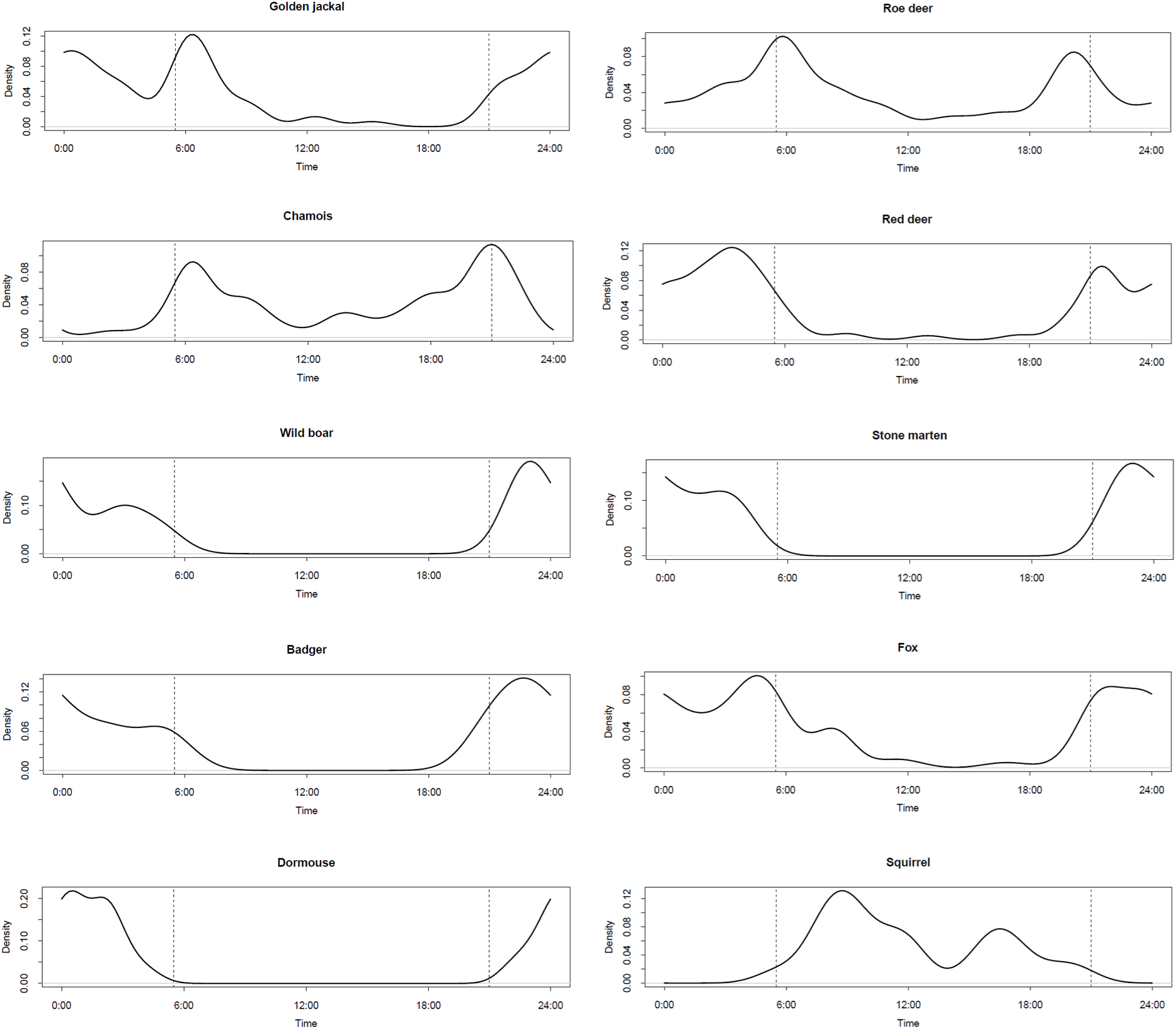
species activity patterns: dashed lines represents dawn and sunshine referred as an intermediate date of sampling period (15 July 2024). Each plot represent a 24h period.

Specifically, golden jackal shows an activity peak around 0:00, followed by a decrease immediately before dawn and a sequent peak after 6:00 am higher than the previous one. The activity remains then low during the day. Roe deer and chamois show crepuscular activity, with negative peaks during the day for roe deer and during the night for chamois. The red deer shows a similar pattern but has higher nocturnal activity. The wild boar has a peak before midnight and scarce activity during the day. Stone marten and badger show nocturnal activity, similar to fox, whose activity is however more dilated during the night and includes dawn. The fat dormouse activity is characterized by a peak in the middle of the night, squirrel is diurnal with a negative peak afternoon.

In the first group of results, we select golden jackal as predator. The Watson two-sample test results are significant for the majority of species (p-value < 0.05), with the exception of fox, badger and red deer fawns. Activity curves of roe deer, red deer adults, chamois, wild boar and stone marten are therefore significantly different from golden jackal ones. The highest overlap coefficient is with fox (Δ_1_ = 0.75) however the test is not significant (p-value >0.051). The same situation occurred with fawns and badger. In table 4 we reported the overlap coefficients for the species with respect to golden jackal, Watson two-sample test values and related significance.

**Table 4:** overlap coefficient calculation results (Δ_1_) of activity patterns with respect to golden jackal. We also report Watson two sample test results (U^2^) with respective significance values.

| Overlap coefficients with respect to golden jackal |  |  |  |
| --- | --- | --- | --- |
| Species | $\Delta_1$ | $U^2$ | p-value |
| *Roe deer* | 0.66 | 0.63 | < 0.001 |
| *Red deer fawns* | 0.64 | 0.17 | 0.05 < p-value < 0.10 |
| *Red deer adults* | 0.71 | 0.25 | < 0.05 |
| *Fox* | 0.75 | 0.17 | 0.05 < p-value < 0.10 |
| *Stone marten* | 0.60 | 0.53 | < 0.001 |
| *Badger* | 0.67 | 0.16 | 0.05 < p-value < 0.10 |
| *Chamois* | 0.61 | 0.36 | < 0.01 |
| *Wild boar* | 0.72 | 0.19 | < 0.05 |

More in detail, the roe deer shows a negative peak of activity when the golden jackal reaches its first positive peak, however, the prey’s second peak is centered around the same hours (Fig. 6a). Fawns and chamois show similar results (Fig. 6b, 6g). On the opposite, red deer adults result in high nocturnal activity with a decrease in correspondence of predator second peak (Fig. 6c). Stone marten is active in the same time interval as predator; however, it presents a slight decrease in activity in correspondence with jackal’s first peak (Fig. 6e). The same can be argued for badger where the decrease is even more pronounced (Fig. 6f). Fox peaks show a weak anticipation with respect to golden jackal ones (Fig. 6d). Finally, wild boar shows a decrease in activity before the predator’s second peak and a slight decrease in correspondence to jackal first activity peak (Fig. 6h).

**Figure 6:**
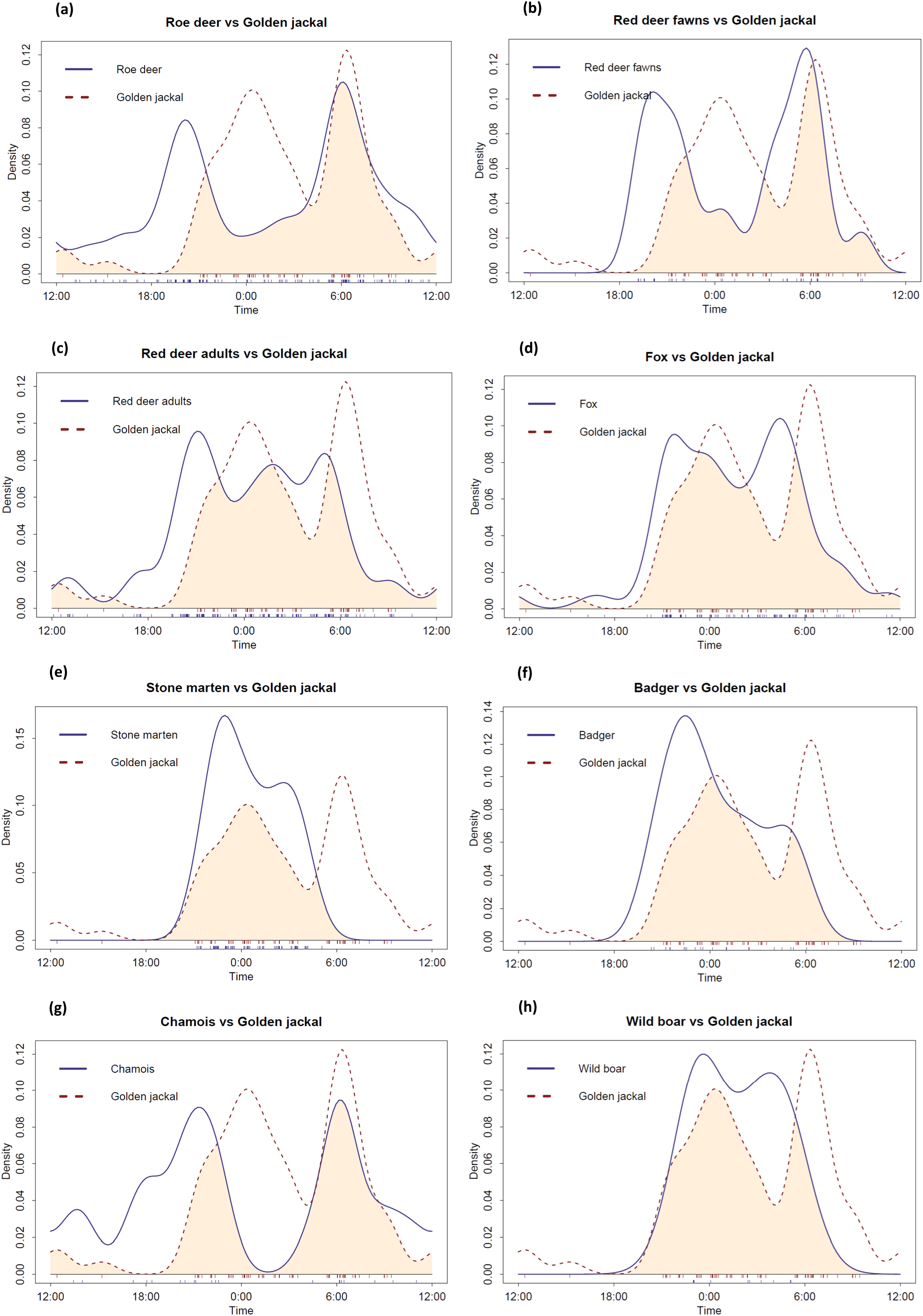
species activity patterns overlap respect to golden jackal: dashed line represents golden jackal activity curve, solid line specie selected. Colored area below both curves it is equal to calculated coefficient. Comparison with a) roe deer, b) red deer fawns, c) red deer adults, d) fox, e) stone marten, f) badger, g) chamois and g) wild boar.

In the second group of results, we select the fox as predator (Table 5). All species considered have activity curves significantly different from fox one. The squirrel shows the lowest overlap values with respect to fox (Δ_1_ = 0.23), followed by dormouse (Δ_1_ = 0.52) and stone marten (Δ_1_ = 0.68). Both stone marten and dormouse concentrate their activities during the night, while instead their predator shows a more dilated curve, with periods of activity even before sunset and after sunrise (Fig. 7a, 7c). There are evident differences in activity patterns with respect to the squirrel (Fig. 7b).

**Figure 7:**
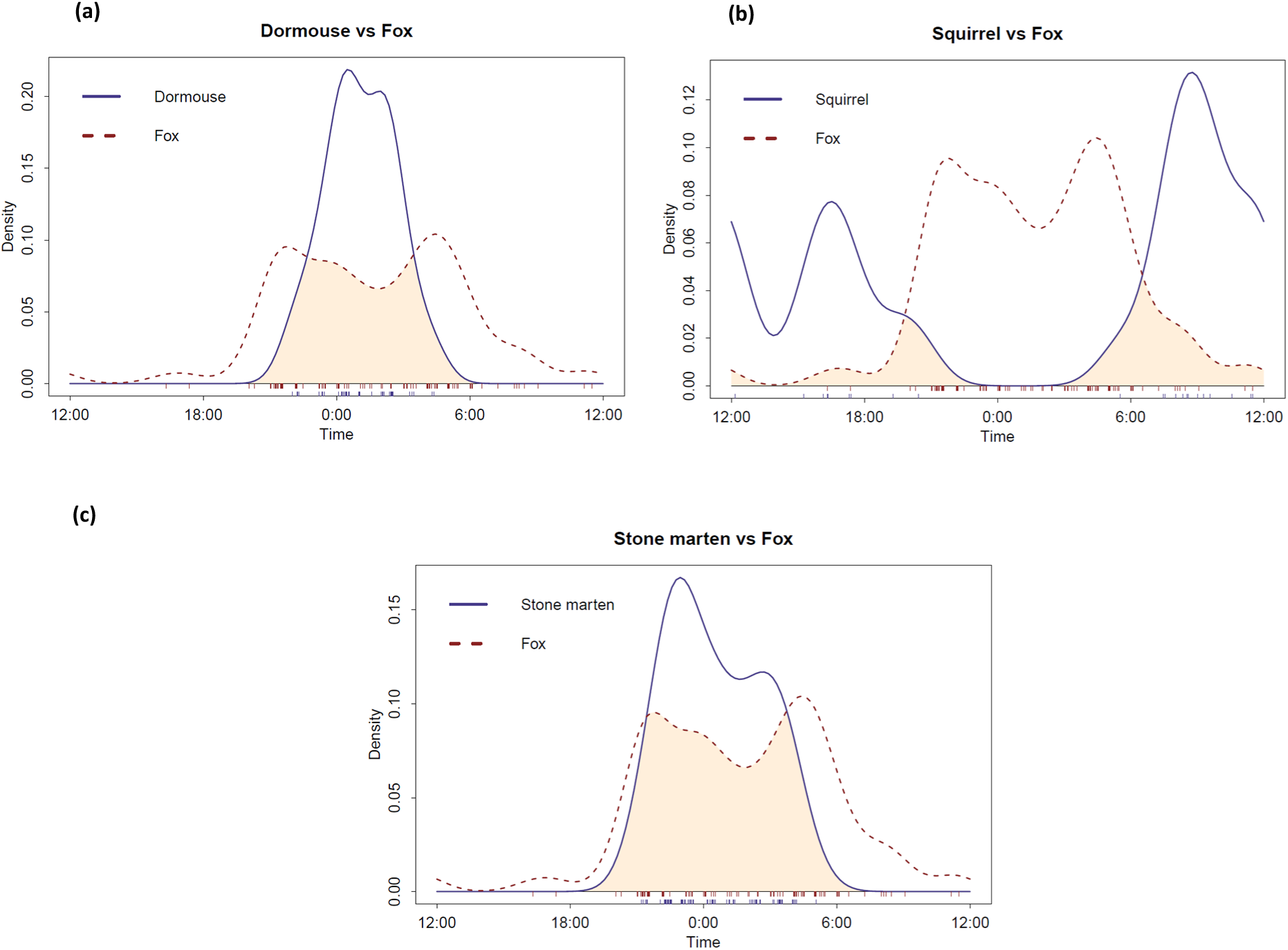
species activity patterns overlap respect to fox: dashed line represents fox activity curve, solid line specie selected. Colored area below both curves it is equal to calculated coefficient. Comparison with a) dormouse, b) squirrel, c) stone marten.

**Table 5:** overlap coefficient results (Δ_1_) of activity patterns of prey with respect to fox. We also report Watson two sample test results (U^2^) with respective significance values.

| Overlap coefficients with respect to fox |  |  |  |
| --- | --- | --- | --- |
| Species | $\Delta_1$ | $U^2$ | p-value |
| *Dormouse* | 0.52 | 1.08 | < 0.001 |
| *Squirrel* | 0.23 | 1.26 | < 0.001 |
| *Stone marten* | 0.68 | 0.72 | < 0.001 |

In the third group of results, we select the stone marten as predator (Table 6). The overlap between the predator and the squirrel is very small and the two distributions are significantly different (p-value < 0.05) (Fig. 8a). The dormouse, instead, shows a high overlap in activity curves, they have both high nocturnal activity (Fig. 8a). However, Watson’s two-sample test is significant (p-value < 0.05).

**Figure 8:**
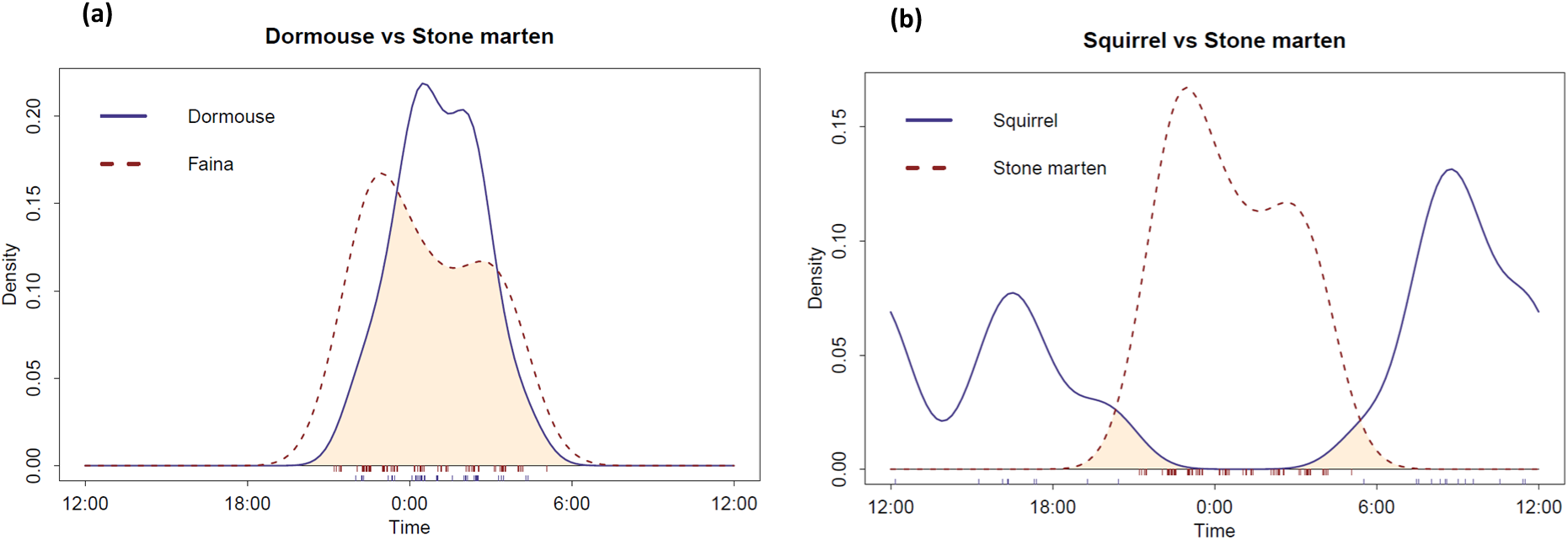
species activity patterns overlap respect to stone marten: dashed line represents stone marten activity curve, solid line specie selected. Colored area below both curves it is equal to calculated coefficient. Comparison with a) dormouse, b) squirrel.

**Table 6:** overlap coefficient calculation results (Δ_1_) of activity patterns with respect to stone marten. We also report Watson two sample test results (U^2^) with respective significance values.

| Overlap coefficients with respect to stone marten |  |  |  |
| --- | --- | --- | --- |
| Species | $\Delta_1$ | $U^2$ | p-value |
| *Squirrel* | 0.07 | 1.54 | $< 0.001$ |
| *Dormouse* | 0.73 | 0.41 | $< 0.001$ |

**Table 7:**
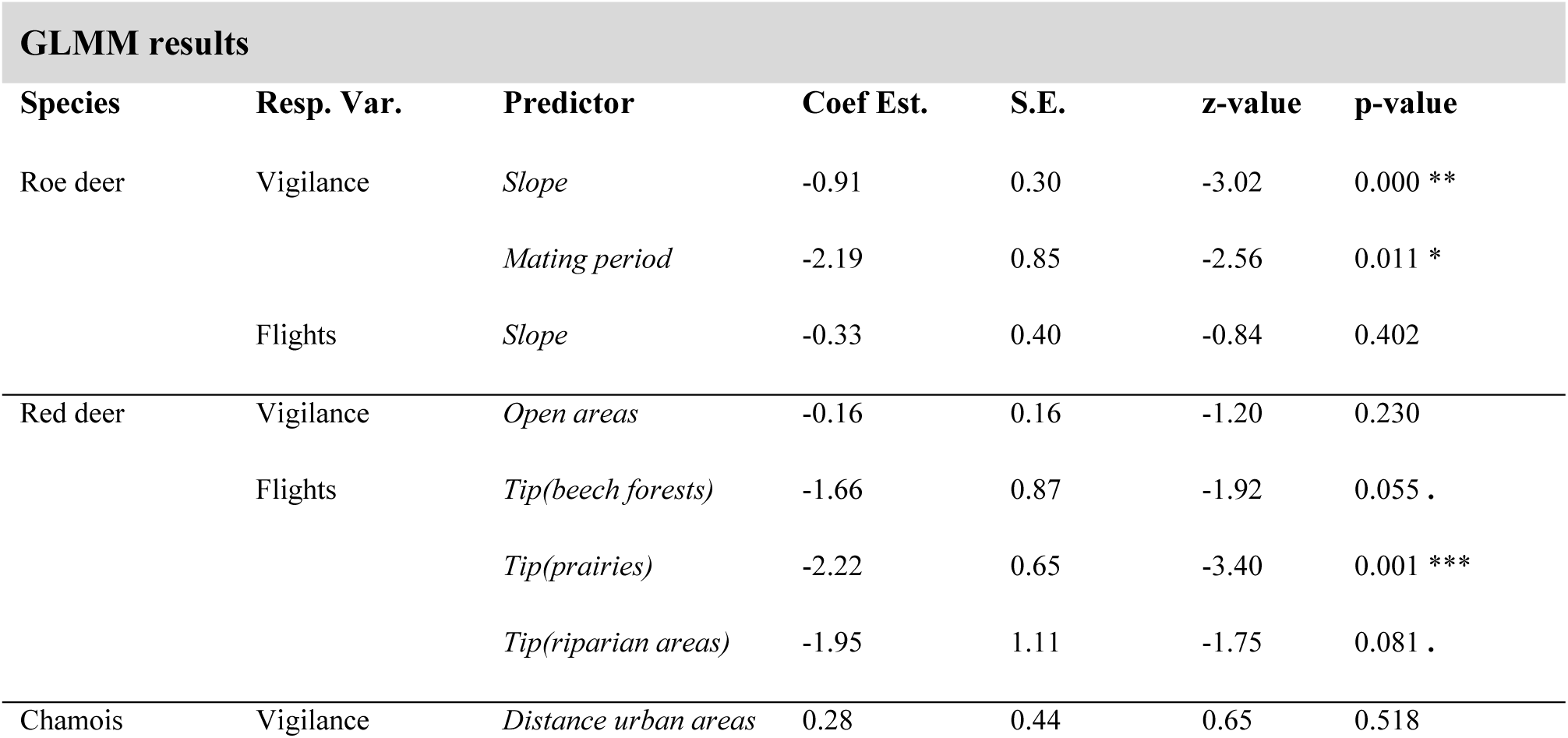
GLMM selected for roe deer and red deer behavioral traits. For each model we report: coefficient estimation (Coef. Est.), standard error (S.E.), statistical test value (z-value) and related significance (p-value)

| GLMM results |  |  |  |  |  |  |
| --- | --- | --- | --- | --- | --- | --- |
| Species | Resp. Var. | Predictor | Coef Est. | S.E. | z-value | p-value |
| Roe deer | Vigilance | *Slope* | -0.91 | 0.30 | -3.02 | 0.000 ** |
|  |  | *Mating period* | -2.19 | 0.85 | -2.56 | 0.011 * |
|  | Flights | *Slope* | -0.33 | 0.40 | -0.84 | 0.402 |
| Red deer | Vigilance | *Open areas* | -0.16 | 0.16 | -1.20 | 0.230 |
|  | Flights | *Tip(beech forests)* | -1.66 | 0.87 | -1.92 | 0.055 . |
|  |  | *Tip(prairies)* | -2.22 | 0.65 | -3.40 | 0.001 *** |
|  |  | *Tip(riparian areas)* | -1.95 | 1.11 | -1.75 | 0.081 . |
| Chamois | Vigilance | *Distance urban areas* | 0.28 | 0.44 | 0.65 | 0.518 |

### Behavioral traits analysis

None of the behavioral traits analyzed for ungulates results related to predator presence probability. Among carnivores, only stone marten interaction with the attractant increases with golden jackal presence probability. The other fox and stone marten traits do not show any relation with predators’ presence probability.

Vigilance, in top ranked model for roe deer, decreases with increasing slope (p-value < 0.001) (Fig. 9a). In addition, also the effect of mating season results significant, with high vigilance values outside this period (p-value < 0.011). The model considering red deer vigilance is not significant (p-value = 0.230), as for chamois (p-value = 0.518). Red deer flights are related to habitat typology with frequent flights in coniferous forests. In this case, roe deer flights model is not significant (Fig. 9b).

**Figure 9:**
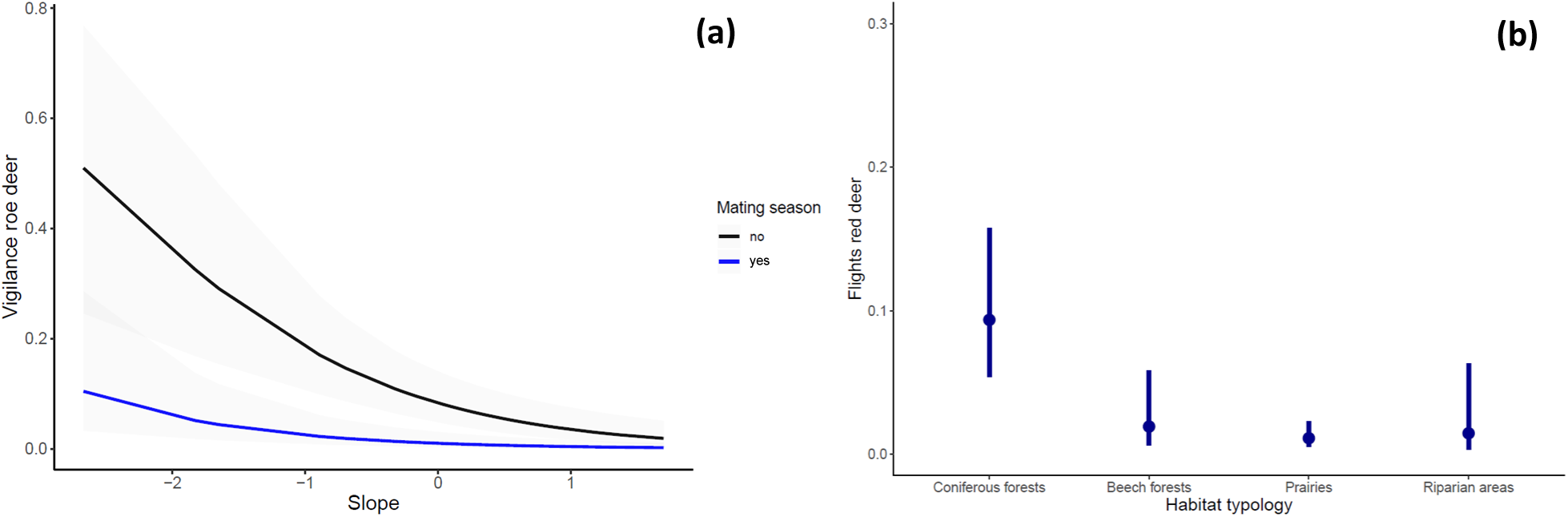
GLMM previsions for a) roe deer: vigilance decreses with increasing slope and it’s higher outside mating season. Colored line represents the relation trend; in light grey 85% confidence interval. Indipendent variables are represented in standardized scale. For b) red deer: highest flight values are in coniferous woods. Dot represents mean calculated value and the line the 85% confidence interval.

Fox indifference to the bait increases with the proportion of forest cover in the landscape (p-value = 0.026) (Fig. 10a), and with altitude (Fig. 10b), but in the latter case, p-value is not significant (p-value = 0.071). Fox interaction with bait increases with altitude (p-value = 0.077) and the latency model is far from significance (p-value = 0.575) (Fig. 10c). Stone marten indifference increases with altitude (p-value = 0.091) (Fig. 11a), interaction with golden jackal presence probability (p-value = 0.083) (Fig. 11b) and latency is greater in beech forests (p-value < 0.001) (Fig. 11c). For golden jackal, none variable results significantly correlated with measured behavioral traits. In table 8 we report top ranking models of each species and related statistics.

**Figure 10:**
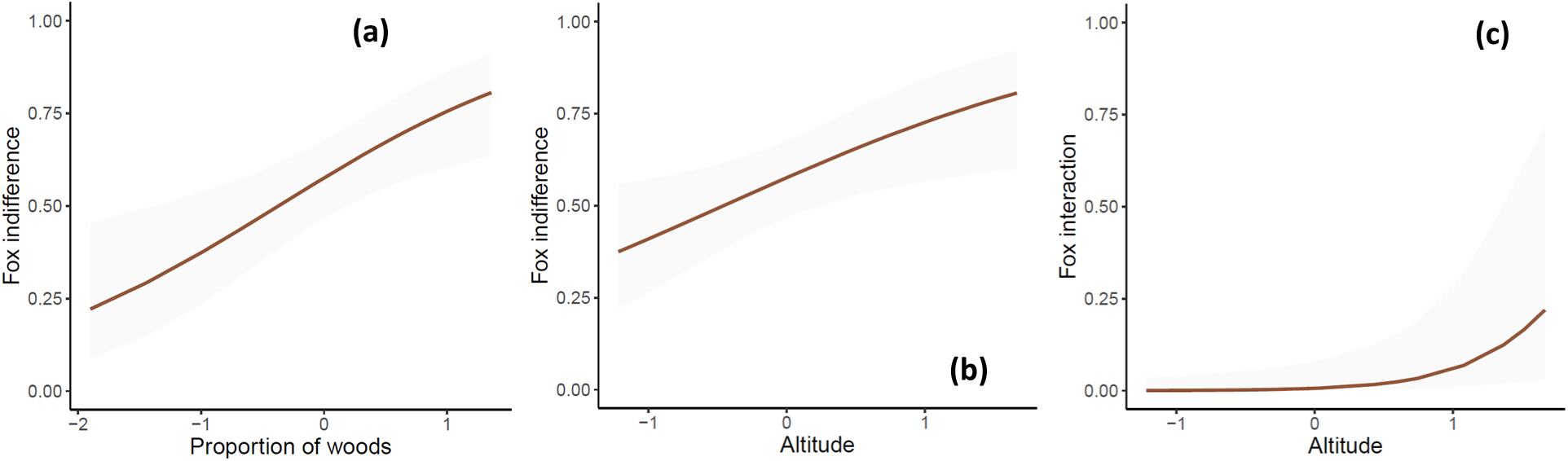
GLMM previsions for fox: indifference to the attractant increases with proportion of woods (a) and with altitude (b). In c) interaction with attractant increases with altitude. Colored lines represent the relation trend; in light grey 85% confidence interval. Indipendent variables are represented in standardized scale

**Figure 11:**
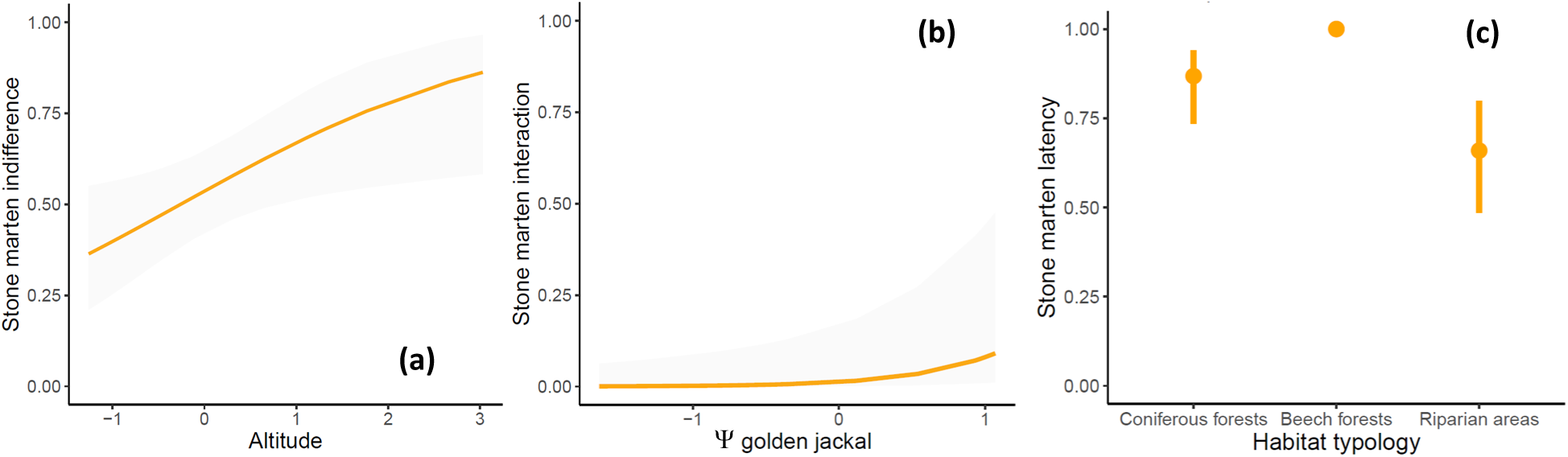
GLMM previsions for stone marten: a) indifference to the attractant increases with altitude; b)interaction with attractant increases with golden jackal presence probability. Colored lines represent the relation trend; in light grey 85% confidence interval. Indipendent variables are represented in standardized scale. In c) latency at the contact with attractant is maximum in beech forests and remains high in the other habiyat typologies; line represents 85% confidence interval.

**Table 8:** GLMM selected for fox, stone marten and golden jackal behavioral traits. For each model we report: coefficient estimation (Coef. Est.), standard error (S.E.), statistical test value (z-value) and related significance (p-value)

| GLMM results |  |  |  |  |  |  |
| --- | --- | --- | --- | --- | --- | --- |
| Species | Resp. Var. | Predictor | Coef. Est. | S.E. | z-value | p-value |
| Fox | Indifference | *Wood* | 0.82 | 0.37 | 2.22 | 0.026 * |
|  |  | *Altitude* | 0.67 | 0.37 | 1.80 | 0.071 . |
|  | Latency | *Altitude* | - 0.84 | 1.50 | - 0.56 | 0.575 |
|  | Interaction | *Altitude* | 2.28 | 1.29 | 1.77 | 0.077 . |
| Stone marten | Indifference | *Altitude* | 0.56 | 0.33 | 1.69 | 0.091 . |
|  | Latency | *Tip(beech forests)* | 19.71 | 0.78 | 25.12 | 0.000 *** |
|  |  | *Tip(riparian areas)* | -1.22 | 0.76 | -1.61 | 0.107 |
|  | Interaction | *Wood* | 3.41 | 2.26 | 1.51 | 0.131 |
|  |  | *Golden jackal* | 1.94 | 1.12 | 1.73 | 0.083 . |
| Golden jackal | Indifference | *Slope* | 0.37 | 0.34 | 1.11 | 0.267 |
|  | Latency | *Distance urban areas* | 4.20 | 3.78 | 1.11 | 0.267 |
|  | Interaction | *Slope* | -1.16 | 1.09 | -1.07 | 0.286 |

## DISCUSSION

To contribute to the understanding of the mechanisms by which animals cope with the landscape of fear, we analyzed together the three main antipredator responses to understand which strategy is chosen by prey to avoid predation risk. We studied, in particular, the influence of predation risk and environmental variables on the habitat selection process, daily activity patterns, and vigilance behaviour. Results show that selection for time intervals where predators are less active is the most widespread antipredator strategy adopted by mammal species studied. On the contrary, results exhibit no responses in terms of habitat selection, except for the fox, which shows a negative relation with golden jackal presence probability; however, this relation looks secondary with respect to the presence of human settlements. Finally, behavioral traits are related more to environmental variables than to predators’ presence probability. Thus, in summary, prey in the studied area respond to the risk of being killed by selecting for temporal intervals in which their predator are less active. We, therefore, demonstrate the existence of a landscape of fear capable of influencing prey decisions in terms of activity periods (at the analyzed scale). The method proposed here has proven to be useful for studying how fear can affect communities and ecosystems.

### Occupancy analysis

None of the top-ranking models included predator presence probability among selected variables. Species seem to prioritize other environmental characteristics in selecting their habitat. Only fox and squirrel include predator presence probability among predictors in selected models; however, these models are not the most supported. For chamois and wild boar, some decisions may be indirectly related to fear of predators, in particular to different habitat characteristics, particularly adapted to different hunting strategies. For all the other species, at the analyzed scale, habitat selection is determined by environmental characteristics.

The selection of a home range for an animal (i.e., second-order habitat selection) is the result of choices that depend on each territory and on relations among different species (e.g., competition, predation) (Johnson, 1980; Rettie and Messier, 2000). Different areas put the animals in front of different choices; in addition, high-quality patches are selected by dominant individuals, excluding the others in low-quality ones. Thus, sometimes, avoiding areas inhabited by predators becomes unfeasible. This could be the case of roe deer: it avoids steep areas due to high energy costs and lack of specific adaptations (Amici et al., 2011; Gaudry et al., 2015), which can be the most important limiting factor in roe deer habitat selection decisions. In other cases, species face different predators and respond to risk at different scales within their home range (i.e., third-order habitat selection). The examples here are the dormouse, whose predators are not only foxes and martens but also owls (Juškaitis, 2023; Kryštufek, 2010), and the stone marten, predated by both foxes and jackals (Hatlauf and Lanszki, 2024; Roy et al., 2019). In addition, the golden jackal is a new predator for the area analyzed; thus, prey, like roe deer, may not be able to detect predator cues and develop efficient antipredator strategies. Also, the tradeoff between safety and resource may lean toward food acquisition: monospecific forest stands may lack sufficient prey even for generalists like the stone marten, leading this species to select for urban or riparian habitats regardless of predation risk (Duduś et al., 2014). Finally, for species like the squirrel, we had only limited data at our disposal, possibly obscuring the relation (stone marten presence probability negatively affects squirrel presence probability, but the model is not the top-ranking one). A similar situation appears between fox and golden jackal (i.e. partial evidence of negative relation between presence probabilities): other studies demonstrate a negative impact of golden jackal on fox due to similar size and ecological niches (Newsome et al., 2017; Tsunoda, 2022; Tsunoda et al., 2018). However, when the resources are abundant (e.g., near human settlements) or the niches only partially overlap (e.g., hunting different prey), the coexistence has been demonstrated (Alexandre et al., 2020; Sévêque et al., 2020). The latter may be our case, considering that the golden jackal selects for areas with a high presence probability of roe deer, suggesting a selection for this prey as reported in other works (Torretta et al., 2021).

In other cases, certain results may hint at innate risk responses even if predators are absent or occasional in the area: chamois detection probability decreases in coniferous forests, where the understory level is well developed. One of its predators is the lynx (*Lynx lynx*), namely an ambush predator, highly efficient in denser patches: during summer, chamois may avoid these riskiest areas thanks to the prairie’s resource abundance, moreover, ambush predators usually boost ungulates’ antipredator responses (Corlatti et al., 2022; Epperly et al., 2021; Wikenros et al., 2015). Similarly, wild boar presence probability decreases with the amount of beech forests in the landscape, which usually has a poor understory, thus a lack of cover (Meriggi and Sacchi, 2001). On the contrary, they show a selection for urban areas as reported in other works (Massei et al., 2015); however, also for this species, we have only a few presence data, as can be seen by the relatively wide confidence intervals.

For the other species, the detection probability of red deer and stone marten decreasing with riparian habitats’ abundance is probably the result of technical constraints in camera traps’ displacement in these rough areas, or, for the latter species, due to higher food availability decreasing the interest for the attractant. Constant presence probability for badger and chamois may be the result of a lack of correct variables capable of explaining variability, such as soil characteristics for the badger (Roca, 2014). Wild boar detection probability increases with beech forest availability, given the high relevance of beech nuts in its diet (Cutini et al., 2013). The most relevant result is probably the positive relation between the golden jackal and the roe deer presence probability. The former is known to be an opportunistic predator shifting its diet according to the abundance of prey, including ungulates where available, even if direct predation is debated and probably focused on fawns (Hayward et al., 2017; Lange et al., 2021; Lanszki et al., 2016; Tsunoda and Saito, 2020). However, here we demonstrate a direct selection for roe deer in the jackal habitat selection process. In addition, these predators tend to avoid high altitudes (Tsunoda, 2022) and to be highly detectable near human settlements, probably due to higher confidence with unusual scents (i.e., attractant). Finally, the fox selects for urban areas where it can easily find different food sources (Alexandre et al., 2020; Contesse et al., 2004; Vuorisalo et al., 2014). Among arboreal rodents, dormouse detectability is higher when there is a high percentage of wood in a radius of 100 m, probably due to the high frequentation of these areas (Juškaitis, 2023; Kryštufek, 2010). The squirrel is more detectable in coniferous forests as the principal source of food; however, presence probability shows no relation with this habitat, probably due to scarce presence data (Kenward et al., 1998; Lurz et al., 1998)

### Overlap analysis

Most prey show activity curves significantly different from their predators, which is in support of the hypothesis that this is the preferred antipredator strategy in this area. This response has been demonstrated in other contexts, usually alternatively to habitat selection (Kohl et al., 2018; Smith et al., 2019), revealing an increase in prey diurnal activities (Kamler et al., 2007; Rossa et al., 2021). However, human activity constrains animals to be active during the night period (Gaynor et al., 2018), making our results more relevant in light of the widespread human use of the area. Among ungulates, roe deer, chamois and red deer fawns show high crepuscular activity usually interpreted as an antipredator response (Kamler et al., 2007; Swinnen et al., 2015), even if Bonnot et al., 2020 demonstrate how an increase in predation risk not necessarily increases roe deer crepuscular activity, suggesting how other factors enter the picture. In addition, chamois is reported as a diurnal species (Thel et al., 2024). However, the three ungulates show high confidence interval overlap, and red deer adults have a high nocturnal activity together with a larger overlap coefficient, possibly suggesting a behavior aimed at avoiding the golden jackal’s peak of activity. The Watson two sample test for fawns is not significant; this may be related to the small sample size, but it shows an interesting trend, possibly consolidated with future data. Further support to the hypothesis is given by the fact that native predators of these ungulates usually hunt during the night in response to human presence (Ciucci et al., 1997; Heurich et al., 2014), and ungulate vision is more efficient at crepuscular hours (D’Angelo et al., 2008). However, due to activity overlap at dawn, the efficacy of this antipredator strategy has to be further analyzed, particularly for roe deer. The wild boar does not show to select for a time when the predator is less active, probably because adults are not hunted by jackals as for the badger (Kostantinov et al.,2022; Lanszki et al., 2016). The fox shows usually a spatial response to golden jackal presence rather than a temporal one, as confirmed by our results (Tsunoda et al., 2018). The stone marten, as a possible prey for both jackals and foxes, shows significantly different activity distribution curve (Roy et al., 2019; Tsunoda et al., 2018). The dormouse, typically nocturnal, shows to differentiate in its activity near the attractive from that of its two predators (i.e., fox and stone marten) (Kryštufek, 2010; Randler and Kalb, 2021). The squirrel is a diurnal arboreal species whose rhythms of activity are different from that of the predators considered.

### Behavioral traits analysis

Results show less support for our hypothesis; traits are mainly influenced by environmental factors. Regarding ungulates, vigilance is an innate behavior that remains even without predator presence (Chitwood et al., 2022; Hunter and Skinner, 1998; Laundré et al., 2001; Le Saout et al., 2015) and that responds to specific cues difficult to consider with camera traps (Creel, 2011; Epperly et al., 2021; Liley and Creel, 2008; Smith et al., 2020). In addition, the risk allocation hypothesis states that, in areas of high risk, prey concentrate on foraging in safer moments (Creel and Christianson, 2008; Lima and Bednekoff, 1999). Here, roe deer vigilance decreases with slope, possibly referring to the risk allocation hypothesis due to roe deer’s inefficient adaptations for these conditions (Donini et al., 2025; Gaudry et al., 2015). However, there is a high correlation between slope and jackal presence probability, possibly indicating slope as a proxy of different factors influencing roe deer vigilance. The red deer have no natural predators in the area; however, flights are lower in prairies where human activity is reduced.

The other traits are not widely used in literature, and the results may be considered a first step in their use as indicators of animal fear. In particular, interaction with the attractant may be considered as a proxy of giving-up density measures (Brown and Kotler, 2004), where less consumption of bait indicates higher levels of fear. Instead, indifference results in a poor indicator of animal fear. Specifically, fox interaction is higher in prairies, habitats avoided by jackals, and less used by humans, possibly reflecting, as predicted, higher confidence by this species in prairie areas. On the contrary, stone marten interaction increases with golden jackal presence probability; however, in a giving-up density experiment Diserens et al., 2022, found similar results for *Nyctereutes procyonoides,* suggesting certain species concentrate food research when they feel safe, even at high risk. For both species, indifference increases with altitude, contrary to our expectations, however, this trait may reflect other environmental characteristics such as food availability. Finally, the latency of the stone marten is higher in beech forests: lack of undergrowth may result in more cautious behavior as the scent of carrion may suggest the presence of a predator. However, our sampling period coincides with the stone marten mating one, possibly influencing animal boldness.

### Limits of our study

We acknowledge that our study has been conducted during a single season of sampling (i.e., summer) and only in one year. Things may change over time, mainly due to resource availability (Kohl et al., 2018; Riginos, 2015) or change among individuals, due to different personalities or body conditions (Bleicher, 2017). In addition, we used an attractant to increase detection probability, which if on one hand increase our statistical power (Buyaskas et al., 2020; Mortelliti et al., 2024) it also may attract individuals from away, inducing potential bias in the habitat selection analysis.

### Conclusions

In conclusion, we demonstrate the presence of a landscape of fear in species of the community analyzed, which mainly influences their temporal activity. We found that considering together the possible antipredator responses is a useful method to have a more accurate knowledge of prey decisions. In addition, we have demonstrated a specific selection by golden jackal for areas where the roe deer is present, suggesting a specialization for this carnivore on this ungulate, possibly favoring coexistence with fox. Future studies will focus on these relations and their consequences to understand the effective impact of the golden jackal in the native communities, as well as further the study of antipredator responses considered together.

## Notes

### Competing Interest Statement

The authors have declared no competing interest.

### Funding

- Funded by European Union-Next Generation EU, Mission 4 Component 1 CUP C13C24000100005

